# Microfiber-Templated Porogel Bioinks Enable Tubular Interfaces and Microvascularization Down to the Building Blocks for 3D Bioprinting

**DOI:** 10.1101/2024.12.04.626229

**Authors:** Yuzhi Guo, Ziyu Wang, Xuening Zhang, Jinghang Li, Liliang Ouyang

## Abstract

Vascularization is key to the biofabrication of large-scale tissues. Despite the progress, there remain some outstanding challenges, such as limited vessel density, difficulty in fabricating microvasculatures, and inhomogeneity of post-seeding cells. Here, we introduce a new form of bioink called microfiber-templated porogel (μFTP) bioink to engineer vasculatures down to the filament building blocks of 3D bioprinted hydrogels. The cell-laden sacrificial microfibers (diameter ranges from 50-150 μm) are embedded in the bioink to template tubular voids and deliver endothelial cells for *in-situ* endothelialization. The inclusion of softening hydrogel microfibers retains the desirable rheological properties of the bioink for extrusion-based bioprinting. Such bioinks can be printed into a well-defined 3D structure with tunable tubular porosities up to 55%. Compared to the conventional bulk bioink counterpart, the μFTP bioink supports the significant growth and spread of endothelial cells either embedded in the matrix or sacrificial fibers, free of the post-cell seeding procedure. Furthermore, the bioprinted scaffolds based on μFTP bioink are seen to significantly promote the in-growth of blood vessels and native tissues *in vivo*. The μFTP bioink approach enables the engineering of tubular bio-interfaces within the building blocks and contributes to the *in-situ* endothelialization of microvasculatures, providing a versatile tool for the construction of customized vascularized tissue models.

## 1. Introduction

The vascular network is a major component of the circulatory system in the human body, responsible for the transportation and distribution of blood/immune cells, nutrients metabolic substances.[1] It is involved in the regulation of numerous tissue and organ development and physiological homeostasis and plays an important role in injury repair and disease development.[2, 3] The engineering of biomimetic vascularized tissues can be used for both *in vitro* tissue modeling and *in vivo* tissue/organ transplantation. However, achieving robust vascularization in 3D engineered tissues remains a significant challenge.

A straightforward approach is to culture endothelial cells (ECs) in a suitable extracellular matrix (ECM) that allows for the self-assembly of ECs into blood-capillary-like vasculatures.[4, 5] The formation of functional vascular networks highly relies on the physiochemical properties of ECM materials and other physical and biochemical conditions, such as medium perfusion and growth factor induction.[6–8] Typical ECM materials for this purpose include fibrin and Matrigel, which, however, are often too soft and less adaptable to advanced biofabrication scenarios, such as 3D bioprinting. Combining ECM materials with other complementary materials, such as gelatin or alginate, can enhance their structural integrity. However, it is still challenging for these ECM materials alone to support complex bioprinted structures. Another approach is to fabricate biomimetic lumen channels first and then seed ECs to form endothelial layers on the inner surface of the channels. Complex biomimetic vascular lumens have been fabricated using extrusion-based,[9, 10] digital light processing[11, 12] and volumetric bioprinting[13] technologies. For example, Grigoryan et al. constructed structures with complex intravascular and multivascular networks using digital light processing based on the optimization of a photoabsorber.[12] However, post-seeding cells into lumens could induce nonuniform distribution of cells and have difficulties in achieving proper orientation for vascular layering. To this end, researchers have used sacrificial inks to deliver ECs on-site while the inks are removed.[9, 10] Nevertheless, due to the resolution limit, conventional bioprinting approaches usually yield vasculatures of hundreds of micrometers, lacking microvasculatures beneficial for surrounding cells. Multiphoton ablation[14, 15] or femtosecond laser irradiation[16] techniques can be used to create finer microvascular structures within cell-laden hydrogels but they are less efficient to scale up.

Templating methods based on salt,[17, 18] oil-water emulsion,[19–21] and sacrificial microgels[22–24] have been commonly used to generate micropores in hydrogels for better mass transfer. With a similar topology to tubular vessels, sacrificial microfibers have been used to induce the generation of microchannels and vascularization in hydrogels. For example, Lee et al. incorporated wetspun poly(N-isopropylacrylamide) fibers with gelatin hydrogels and removed the fibers afterwards for *in vivo* angiogenesis induction.[25] Other studies have used sacrificial hydrogel fibers to template microchannel for *in vitro* tissue vascularization, mainly based on the post-seeding of endothelial cells into the channels.[26] Recently, Lammers et al. demonstrated rapid tissue perfusion by incorporating alginate microfibers into fibrin hydrogels.[27] However, the existing fiber-templating work either relies on the post-cell-seeding procedure or migration of surrounding ECs, making it less controllable for endothelialization, and none of them has been adapted to 3D bioprinting systems.

In this study, we developed a microfiber-templated porogel (μFTP) bioink system for 3D bioprinting and in-situ endothelialization, paving the way for the vascularization of customized tissue biofabrication. We first fabricated alginate microfibers with tunable diameters (50um-150um), which were capable of encapsulating living cells. The μFTP bioink was prepared by supplementing photocrosslinkable hydrogel precursor solution with dispersible alginate microfibers, which was only possible by softening the microfibers beforehand (**Figure 1A**). After bioprinting and photocrosslinking for structure stabilization, the embedded microfibers can be readily dissolved and gently washed out by incubation with low-concentration sodium citrate solution, generating microchannels in bioprinted structure with tunable porosities (up to 55%). The rheology tests and bioprinting practice show that the μFTP bioink has good printability and structural stability, where the embedded microfibers appear to be partially aligned along the printed filaments. After the dissolution of microfibers, the endothelial cells were released from the fibers and attached to the inner wall of the generated channels, forming a confluent endothelial layer (**Figure 1B**). Moreover, subcutaneous implantation experiments confirmed the significant promotion of the ingrowth of blood vessels and native tissues with the μFTP-based bioprints (**Figure 1C**). Collectively, we propose a versatile bioink strategy for constructing tubular lumen voids within the building block level of bioprinted tissues, which enables *in situ* endothelialization inside the lumens, providing a practical solution for the engineering of customized vascularized tissues.

**Figure 1.**
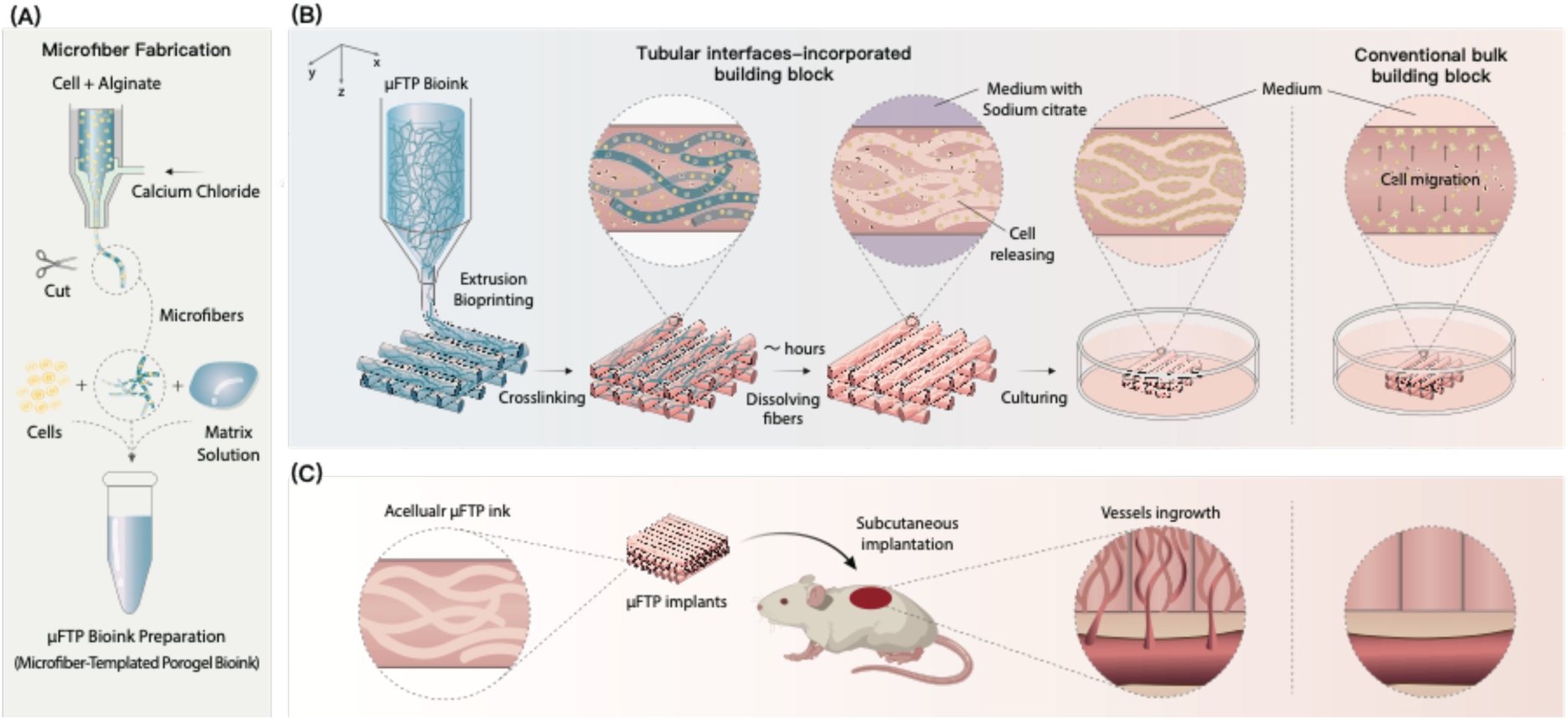
Schematic of formulating μFTP bioink for bioprinting applications. (A) Fabrication of cell-laden microfibers by coaxial microfluidic method and preparation of μFTP bioink. (B) Extrusion bioprinting using μFTP bioink and cell growth process in comparison with conventional bulk bioink counterpart. (C) Schematic of *in vivo* subcutaneous implantation of μFTP scaffolds.

## 2. Results and Discussion

### 2.1. Fabrication of hydrogel microfibers with tunable size and dissociation properties

Due to the quick ionic chelated reaction with divalent cation, alginate has been widely used to fabricate hydrogel microfibers.[28, 29] Here, we use a versatile coaxial flow method to fabricate alginate microfibers with tunable size. 0.5% (w/v) alginate solution and 180 mM calcium chloride solution were respectively injected into the inner and outer layers of a homemade coaxial needle (34G/22G), which is connected to a silicone tube as an outlet for flow restriction. By adjusting the flow rate of the inner and outer layers, continuous alginate microfibers with diameters of 50, 100, and 150 µm were successfully fabricated with good stability and uniformity (**Figure 2A and B**). Decreasing the flow rate of sodium alginate solution in the inner layer would induce significantly smaller microfibers. In contrast, without the outer flow restriction, much larger fibers (e.g., 200 µm in diameter) will be obtained using the 34G needle alone. The fabricated alginate microfibers could be stored in a calcium chloride solution for a long term.

**Figure 2.**
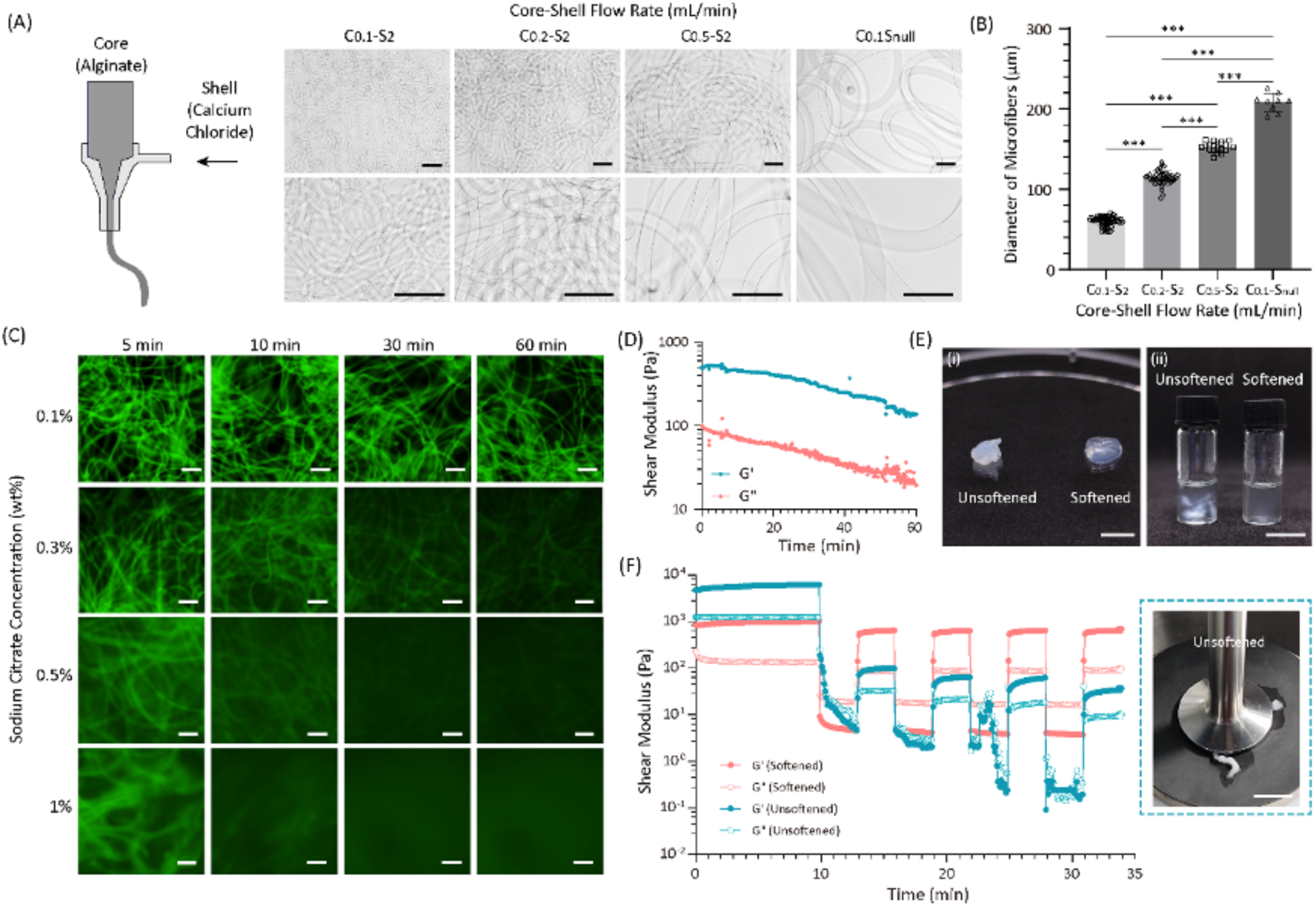
Fabrication and dissolution of tunable hydrogel microfibers. (A) Representative microscopic images of alginate microfibers and (B) the quantified diameters under varying flow rates of inner (0.5 % alginate) and outer (180 mM calcium chloride) layer in coaxial needle. (C) Representative fluorescence images of alginate fibers treated with sodium citrate at different concentrations, illustrating the microfiber’s dissolution profiles. (D) Oscillatory rheological time sweep of collected alginate fibers (diameter of 50 μm) at 22 ℃, frequency of 1.5 Hz and shear strain of 1%. (E) (i) Appearance of packed unsoftened and softened alginate microfibers and (ii) μFTP bioink prepared with unsoftened and softened alginate microfibers. (F) High-low strain cycle tests of unsoftened and softened alginate microfibers and representative image of intertwined untreated microfibers under high strain conditions. Scale bars: 500 μm (A, C), 5 mm (E-i) and 10mm (E-ii, F). One-way ANOVA with a Tukey’s multiple comparisons test, *P < 0.05; **P < 0.01; ***P < 0.001; ns, not significant (n ≥ 9).

Sodium citrate, whose citrate ion can chelate to calcium ions and form calcium citrate complexes, was proven to be an effective chemical to dissolve calcium alginate hydrogels.[30, 31] Here, we used sodium citrate solution with gradient concentrations (0.1%, 0.3%, 0.5%, and 1% (w/v)) to dissolve the alginate microfibers and tested the dissolution profiles. To visualize the dissolution process, alginate was labeled with fluorescein isothiocyanate (FITC) (**Figure 2C**). 0.1% sodium citrate solution is not strong enough to dissociate the fibers in a timescale of 60 minutes. When the concentration of sodium citrate increases to 0.3% or above, the outline of the fiber starts to blur within minutes, and the fluorescence intensity decreases with time, suggesting successful dissolution. As expected, a higher concentration of sodium citrate results in a higher dissolution speed. When using 1% sodium citrate solution, the microfibers were fully dissolved within 30 minutes, while it took approximately 240 minutes to achieve a similar dissolution using 0.3% sodium citrate solution (**Figure S1A**). The rheological tests carried out with pure alginate microfibers indicated the decrease of shear moduli with time treated with sodium citrate solution (**Figure 2D**). This decrease is likely attributed to the dissolution of the microfibers (**Figure S1A**).

We further tested the influences of sodium citrate solution on the growth of endothelial cells. Human umbilical vein endothelial cells (HUVECs) were treated with culture medium supplemented with sodium citrate of varied concentrations for 24 hours (**Figure S1B**). According to the Live/Dead^TM^ staining assay, cell viability is negligibly influenced, but the cellular morphology seems to change with the treatment of high-concentration sodium citrate. The circularity of the cells increased with the concentration increase of sodium citrate, indicating a less adherent tendency of cells when treated with high-concentration sodium citrate. Furthermore, the gene expressions associated with apoptosis (Caspase-3, Bcl-2, and BAX) showed no significant differences between the control and sodium citrate-treated groups with up to 0.6% sodium citrate, indicating the biocompatibility of the treatment process (**Figure S2**). Considering both the dissolution timeline and cell behavior, we chose the 0.3% sodium citrate solution to dissolve alginate microfibers in the subsequent studies.

When manipulating the generated alginate microfibers, we found it difficult to separate them due to the entanglement and sticking, which affected the uniformity of fiber distribution. To better disperse the microfibers in the μFTP formulation, we cut the microfibers into small segments with the length distribution centered from 1.5-3 mm (**Figure S1C**) and softened the microfibers by immersion in saline. The use of a specific volume of saline (20 times the volume of the microfiber precursor solution) would soften the microfibers and make them easier to disperse, resulting in better uniformity of the formulated μFTP bioink. Firstly, saline-treated microfiber has a higher transparency in appearance (**Figure 2E-i**). After pulling the microfibers, we observed that the unsoftened microfibers exhibited irreversible deformation with a reduction in diameter and adhesion between microfibers (**Figure S3**). In contrast, the fractured surface of the softened microfibers appeared smooth, with no significant change in shape (**Figure S3**). In the high-low strain rheological tests, unsoftened microfibers became intertwined under high-strain conditions, which led to inaccurate testing results (**Figure 2F**). In contrast, saline-treated fibers showed lower modulus along with reversible shear-thinning and self-healing properties (**Figure 2F**). This may be because saline washed away excess calcium ions from the surface of the microfibers, making it easier to slip between the microfibers. When preparing the μFTP bioinks, the softened microfibers led to a homogeneous clear formulation, while the inclusion of unsoftened microfibers resulted in unwanted aggregates and precipitations (**Figure 2E-ii**). Together, these results highlight the advantages of softening treatment for microfibers.

### 2.2. μFTP allows for the creation of fibrous pores with tunable porosity

To prepare μFTP bioink, alginate microfibers were cut, softened, and centrifugated to remove the liquid with a 40μm cell strainer. Subsequently, the processed microfibers were mixed with a hydrogel precursor solution, i.e., 7.5% gelatin methacryloyl (GelMA) solution. GelMA was selected as the continuous matrix phase because of its good biocompatibility and easy adaptability for biofabrication. The mixture was treated with light to photocrosslink the matrix and then treated with sodium citrate solution to remove the microfibers. To track the flow of dissociated alginate, we pre-loaded some fluorescent particles (5 μm in diameter) into alginate microfibers. After the treatment with sodium citrate solution, we observed that the fluorescent particles flowed out from the hydrogel structure, which evidenced the successful dissolution and removal of alginate (**Figure 3A**). Compression tests of photocrosslinked μFTP formulation before and after the sodium citrate dissolution process were carried out to further characterize the mechanical properties (**Figure S4**). The results showed that the inclusion of microfibers into the matrix led to a decrease in compressive modulus and an increased tendency for fracturing compared to the bulk hydrogel. After dissociating the microfibers, the compressive modulus of the μFTP hydrogel further decreased to approximately ∼ 3.52 kPa, representing only 66% of its value prior to dissociation.

**Figure 3.**
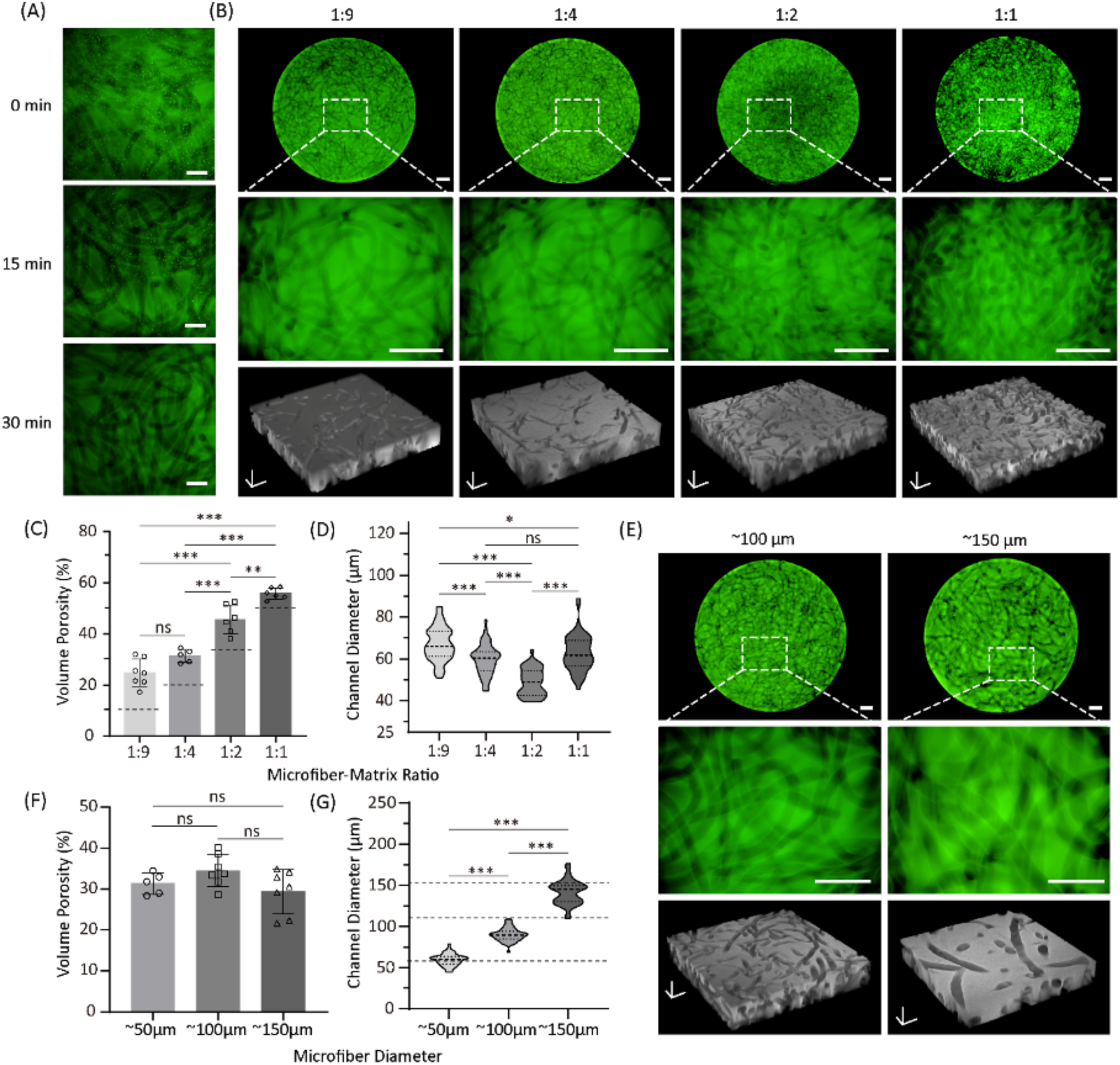
Characterization of microfiber-templated porogels. (A) Representative images of μFTP samples at different time points during 1% sodium citrate treatment, suggesting the dissolution process of microfibers from the matrix. FITC-labeled microparticles are preloaded in the microfiber to indicate the dissolution and location of alginate. (B) Representative fluorescent images and z-stack 3D reconstruction of porogel samples. (C) The measured porosities and (D) pore diameters based on z-stack images under different microfiber to matrix ratios (1:9, 1:4, 1:2, and 1:1). (E) Representative fluorescent images and z-stack 3D reconstruction of μFTP samples. (F) The measured porosities and (G) pore diameters based on z-stack images under different microfiber diameters (50, 100 and 150 μm). The dotted line in C and G indicates the theoretical value of porosity and pore size. Scale bars: 200 μm (A), 500 μm (upper two rows of B and E) and 200 μm (bottom row of B and E in axis). One-way ANOVA with a Tukey’s multiple comparisons test, *P < 0.05; **P < 0.01; ***P < 0.001; ns, not significant (n ≥ 5).

To visualize the porous hydrogel structure and quantify the pores, FITC-labeled GelMA was used as the matrix, and z-stack confocal microscopic images were used to determine the actual porosity. Porous structures using μFTP formulations with varied microfiber-matrix ratios (from 1:9 to 1:1) have been successfully fabricated, with all groups maintaining good structural stability (**Figure 3B**). The microchannels templated by dissolved microfibers were seen to distribute evenly inside the matrix and connect with each other, forming a network with tunable densities. The increase of the initial microfiber fractions contributed to the increase of the eventual porosities, ranging from (24.76±5.38%) to (55.73±2.21%), which were similar to the density of blood vessels in the human body (10% to 50%) (**Figure 3C**). [32, 33] The slight differences between the measured porosities and the theoretical values determined by the fiber fraction might be caused by the remaining water after centrifugation and the swelling behavior of microfibers after mixing with the matrix. Nevertheless, the μFTP formulaton allows us to readily control the porosity of the resulting porous hydrogels. The diameters of microchannels forming by dissolved fibers were also measured to prove the control of pore size (**Figure 3D**).

When using microfibers of 50 µm in diameter, the generated pores exhibit a concentrated size distribution from 40-80 µm throughout the groups with different microfiber fractions. The slight differences in diameter seen between groups might stem from microfiber batch-to-batch variation. We further experimented with microfibers of different diameters to create porogels (100 and 150 µm with a microfiber-matrix ratio of 1:4, **Figure 3E-G**). The porogels presented with accordingly tubular pores are successfully fabricated with excellent stability (**Figure 3E**). There was no significant difference in porosity among groups using microfibers of varied sizes when fixing the microfiber fraction (**Figure 3F**). The further measurements showed a high degree of size consistency between the microchannels and the microfibers (**Figure 3G**). Together, these results demonstrated that our μFTP approach allows for easy control over the tubular porosity and size.

In recent years, microgels have been used as porogens to prepare porous hydrogels due to their high biocompatibility.[22–24] Compared to the microgel-templated method, our μFTP strategy provides a measure to guide the vascular network to be tubular shape, which is more similar to blood vessels than spherical geometry. Furthermore, it allows for better lumen connectivity due to the intrinsic topological differences between spheroids and fibers. To demonstrate the advantages of microfibers over microgels in generating interconnected interfaces, we compared the pore connectivity by using microfibers and microgels with the same volume fraction in the composite ink formulation (**Figure 4A**). Sacrificial alginate microgels were mixed with red fluorescent-labeled GelMA in the same manner as alginate microfibers to prepare hydrogel samples. After dissolving the porogen with sodium citrate solution, the samples were immersed in FITC-labeled dextran for 30 minutes, then immediately imaged using a confocal microscope, followed by 3D reconstruction of the structures using Imaris software (**Figure 4B**). The volumes of the red and green fluorescent regions were quantitatively measured, and the ratio of the green fluorescent volume to the combined volume of the green fluorescent and non-fluorescent regions was used as an indicator of pore connectivity (**Figure 4C**). With the increase of the porogen-matrix ratio from 1:9 to 1:2, both groups exhibited improved pore connectivity, which is from 14.1% to 31.8% in the microgel group and from 28.3% to 55.2% in the microfiber group. At equivalent porogen-matrix ratio, the pores of the μFTP formulation exhibited more extensive dextran infiltration, demonstrating superior pore connectivity in the μFTP system compared with the use of microgel.

**Figure 4.**
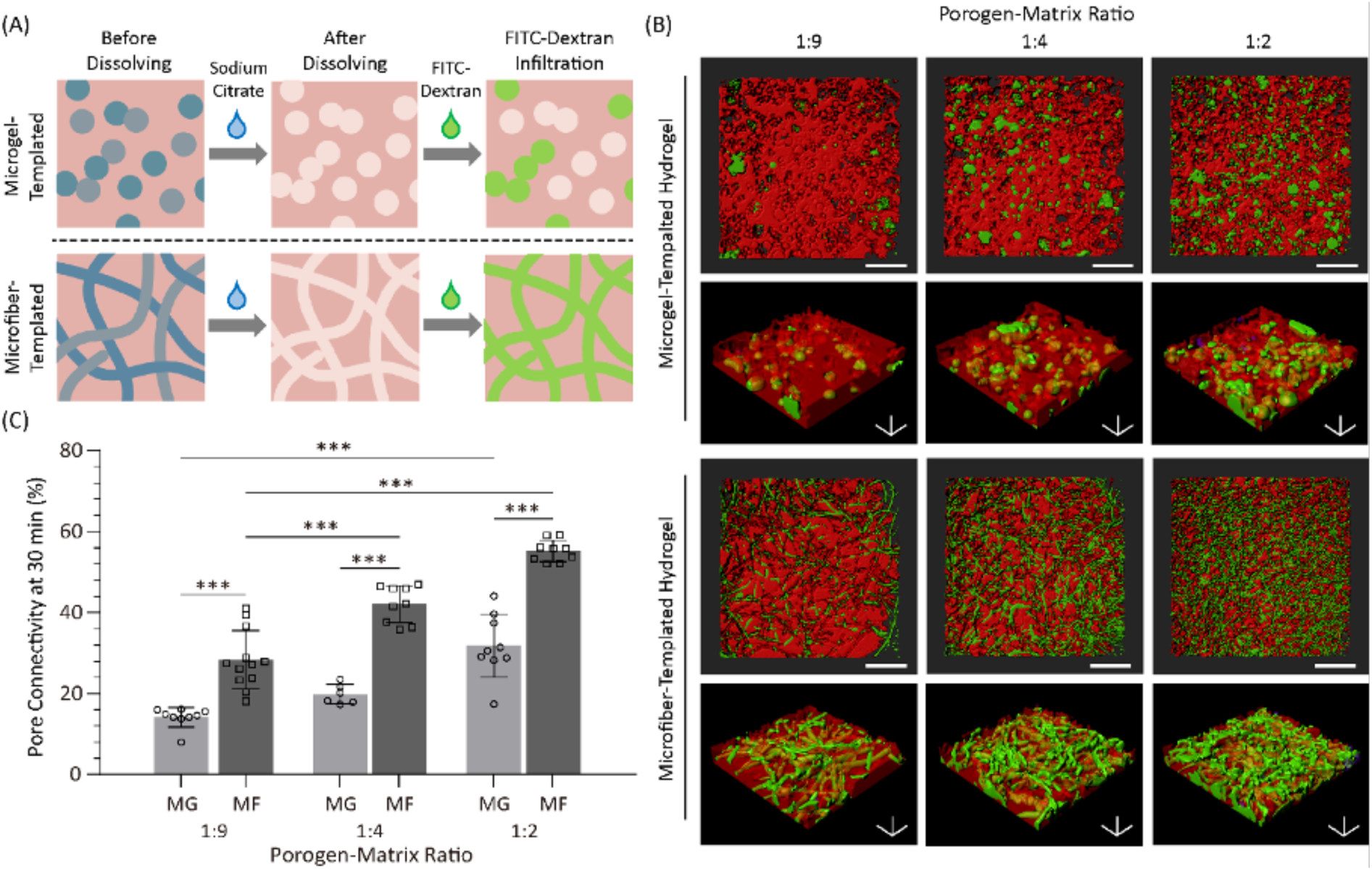
Pore connectivity characterization of microgel- and microfiber-templated porogels. (A) Schematic of the perfusion process of microgel- and microfiber-templated porogel with FITC-dextran. (B) Representative z-stack 3D reconstruction of porous hydrogel structure (red) and infiltrated dextran (green) in which microgels and microfibers serve as porogens at different porogen-matrix ratios. (C) Quantified pore connectivity of porous hydrogel sample with spherical and microchannel pores at different porogen-matrix ratios. Scale bars: 1000 μm (upper row of B) and 200 μm (bottom row of B). One-way ANOVA with a Tukey’s multiple comparisons test, *P < 0.05; **P < 0.01; ***P < 0.001; ns, not significant (n ≥ 3).

### 2.3. 3D printing of μFTP bioinks

To evaluate the printability of the μFTP bioink, we first test its rheological properties. The temperature sweeps indicated that the initial μFTP formulations retain a sol-gel transition property and the transition temperature generally decreased with the increase of microfiber fraction. In general, the sol-gel transition temperatures when cooling are lower than the solutioning temperatures when heating, which coordinates well with the properties of plain gelatin or GelMA formulations without microfibers (**Figure 5A**). Increasing the microfiber fraction will induce a slightly lower sol-gel transition temperature. For example, the gelation temperatures of the μFTP formulations at 1:9 and 1:1 ratios were 26.8 ℃ and 25.2 ℃, respectively. The slight attenuation of the thermo-sensitivity may be directly attributed to the reduced proportion of temperature-sensitive matrix (i.e., GelMA). In addition, the introduction of non-thermo-sensitive microfibers seemed to lead to the weakening of the gels at low temperatures (e.g., 4℃) and strengthing of the sols at high temperatures (e.g., 37℃). For example, the storage moduli at 4 ℃ post cooling decreased from ∼2300 Pa to ∼1300 Pa when increasing the microfiber-matrix ratios from 1:9 to 1:1.

**Figure 5.**
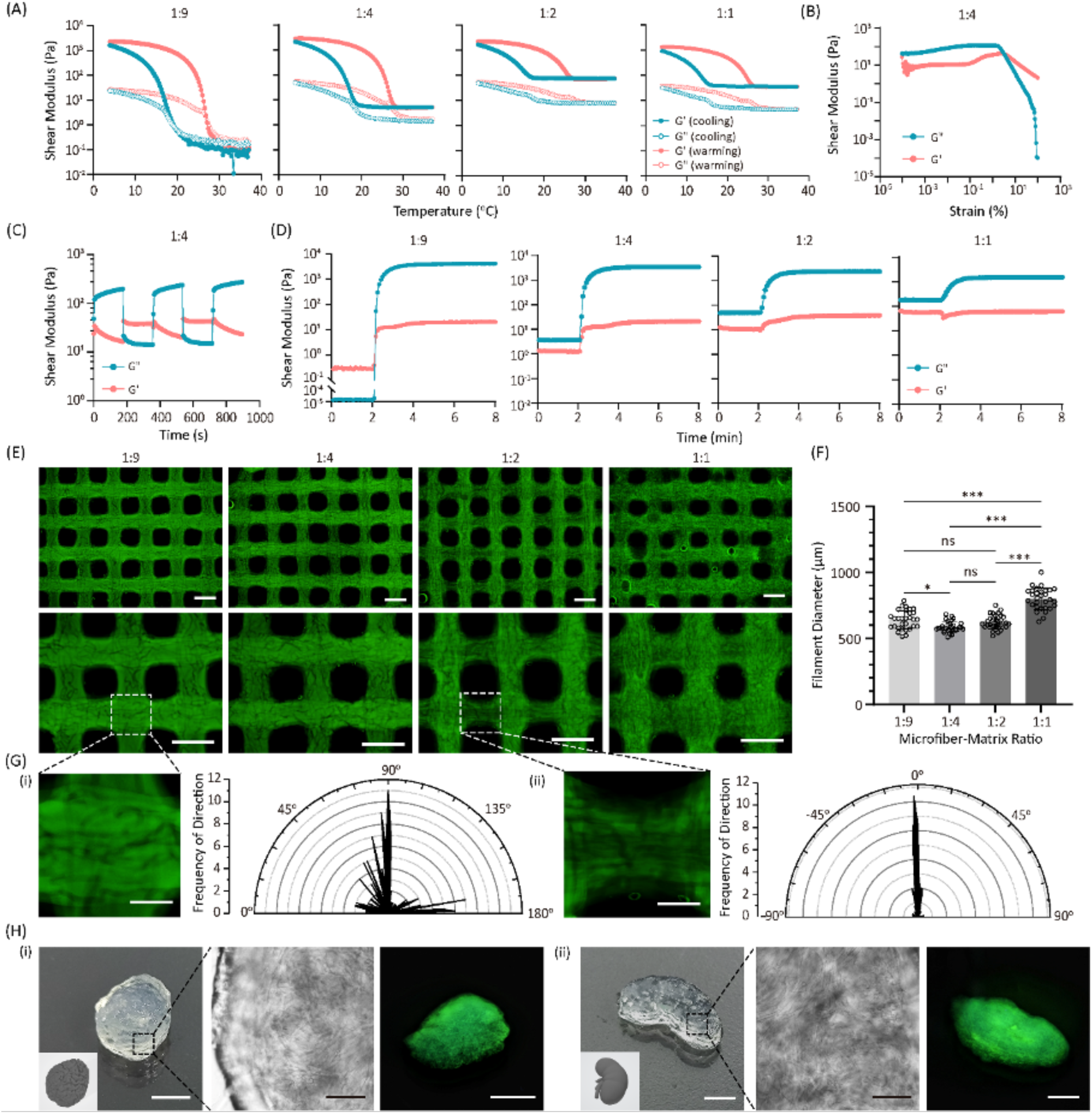
3D printing of microfiber-templated porogel bioink. (A) Temperature sweeps of μFTP bioinks at different microfiber-matrix ratios (1:9, 1:4, 1:2 and 1:1). (B) Strain sweep of μFTP bioinks at 1:4 component ratio. (C) Photocrosslinking rheological tests of μFTP bioinks at different component ratios. (D) High-low strain cycle tests of μFTP bioinks at 1:4 component ratio. (E) Representative microscopic images of printed multilayered lattice structures with different porosities. (F) Quantified filament diameter of printed lattice structure with μFTP bioinks at different component ratios. (G) Representative magnification images of printed scaffolds and the measurements of microfiber orientation in printed filaments. (H) Examples of large-scale samples (i.e., human brain- and kidney-like models) printed using μFTP bioinks at 1:4 component ratio and FITC-labeled dextran were used to test the pore connectivity. All the μFTP bioinks here are formulated using 7.5 wt% GelMA as the matrix. Scale bars: 500 μm (E-F), 200 μm (G), 5 mm (optical and fluorescent photography in H) and 1 mm (microscopy in H). One-way ANOVA with a Tukey’s multiple comparisons test, *P < 0.05; **P < 0.01; ***P < 0.001; ns, not significant (n=30).

Subsequently, strain sweeps were performed on different ink components to further verify the rheological properties of the bioinks (**Figure 5B** and **Figure S5A**). The results showed that all μFTP bioinks behaved like an elastic-like solid at low strains but yields at high strains. The yield point showed a slightly leftward shift as the proportion of microfibers increased, with the yield strain value decreasing from 3.71% to 2.94% when increasing the microfiber ratios from 1:9 to 1:1. A higher microfiber fraction might induce a higher slip tendency, thus leading to a lower critical value of yield strain. Further, the high-low stain cycle tests showed that all the μFTP groups showed a rapid self-healing property (**Figure 5C** and **Figure S5B**). Finally, the photorheological tests indicated that different ink formulations showed similar photocrosslinking kinetics and time profiles, but the μFTP bioinks containing more microfibers presented a lower modulus after complete crosslinking (**Figure 5D**). Overall, the μFTP bioink exhibits excellent shear-thinning and self-healing properties, suggesting the potential to be applied in extrusion-based 3D bioprinting.

Then the μFTP bioinks with different microfiber (50 μm in diameter) fractions were printed into lattice structures under optimized temperature (22 ℃), in which FITC-labeled GelMA (7.5% (w/v)) served as the matrix phase. Under the same printing conditions, there was no significant difference in the diameter of the printed filament among groups with microfiber-matrix ratios from 1:9 to 1:2 (**Figure 5E-F**). When increasing the microfiber fraction to 50%, the filaments were slightly bigger than others (**Figure 5F**), which was likely due to the decrease in modulus (**Figure S5**). Nevertheless, these results revealed the good printability and stability of μFTP with varied microfiber fractions. In addition, the microscopic images show that microfibers oriented to some extent within the lattice structure, which might be due to the shearing and directional alignment during extrusion. We further quantified the orientation of the microfibers (**Figure 5G**). The results showed that the orientations of embedded microfibers were well concentrated at 0° and 90° at the nodes of the lattice structure (Figure 4G(i)), while the orientations of the microfibers were concentrated at 0° in the printed horizontal filament. μFTP bioinks containing microfibers of different diameters (100 μm and 150 μm) with different nozzles (18G, 22G and 25G needles) were also printed (**Figure S6A, B**). It was observed that the diameter of the microfiber had no significant effects on 3D printability. With smaller needles (22G and 25G), fewer microfibers were observed in the printed structure, which might be because fibers would be stuck to the outlet of a thin needle. We also demonstrated the large-scale structural printability of μFTP ink by printing centimeter-scale models such as brain and kidney and treated them with sodium citrate solution afterwards, both of which showed excellent shape fidelity (**Figure 5H and Figure S6C**). The microscopic images confirmed the generation of microchannels and a clear channel orientation along the print path. The large models were also immersed in FITC-labeled dextran, and under UV light, dextran infiltration into the internal microchannels was clearly observed after 12 hours.

### 2.4. In-situ endothelialization of bioprinted structure using μFTP bioinks

To demonstrate the feasibility of in-situ delivery and endothelialization of ECs using the μFTP bioink, we first tested the biocompatibility of the microfiber fabrication process (**Figure 6A** and **Figure S7**). HUVEC-laden alginate microfibers (50, 100, and 150 μm in diameter) can be successfully fabricated with an initial cell density of 2.5×10^6^/mL, and cell viability kept high within 12 hours after fiber fabrication (∼85% after 2 hours and ∼75% after 12 hours) (**Figure 6B**). A longer period of encapsulation in alginate would result in further cell death, likely due to the nonsuitable matrix environment of alginate for the 3D culture of HUVECs. In our protocols, samples were treated with sodium citrate for up to 12 hours, which was sufficient to fully dissolve the alginate and keep the cells alive after being released. Two other endothelial cell lines, hCMEC/D3 (a normally used brain microvascular endothelial cell line) and MS1 (a murine pancreatic islet endothelial cell line) were also used to validate the applicability of our approach. Both types of cells presented viabilities higher than 75% within 12 hours (**Figure S7**), indicating the potential to formulate μFTP bioinks.

**Figure 6.**
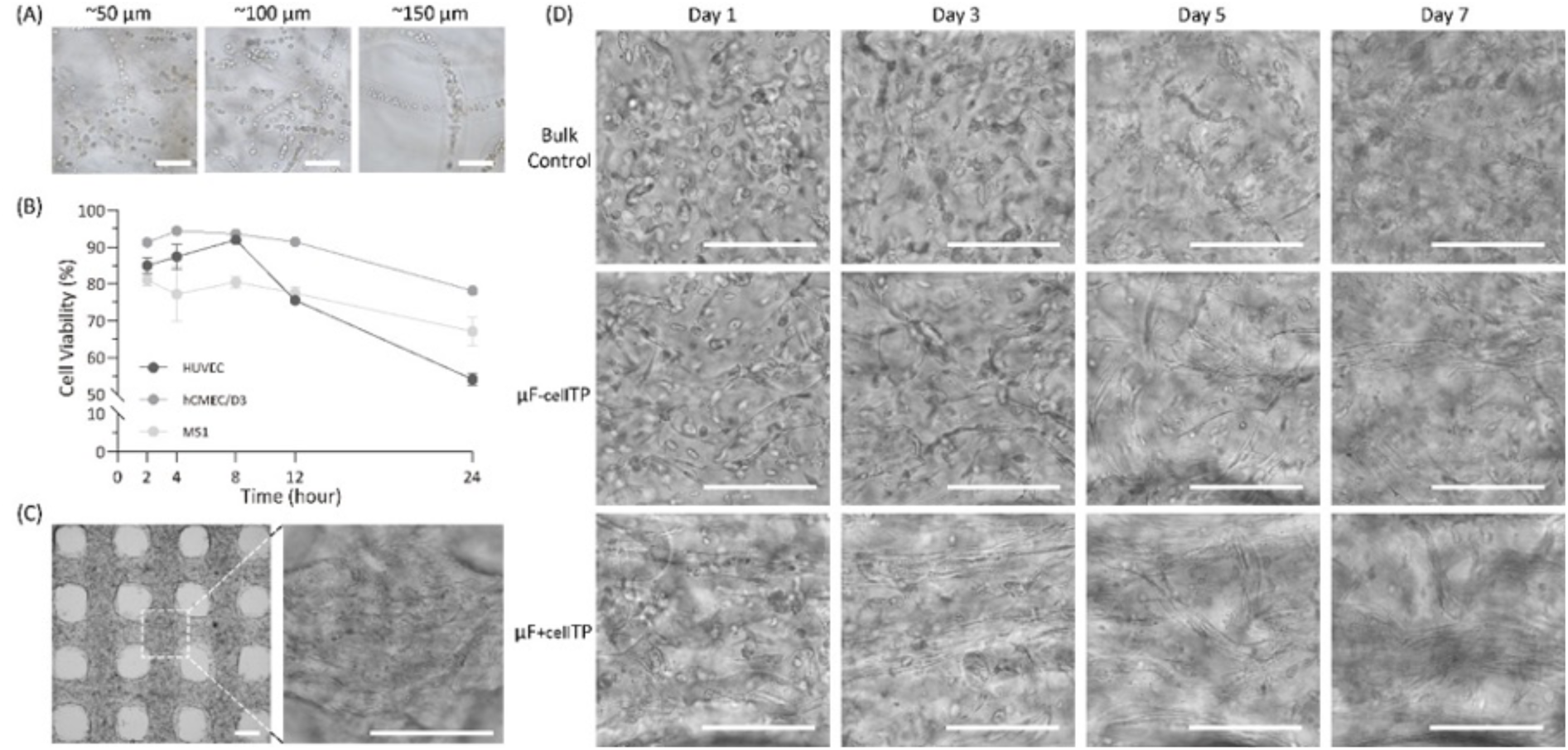
Microfiber-templated porogel bioink allows for spread of HUVECs within 3D printed structures. (A) Microscopic images of fabricated cell-loaded alginate microfibers with different diameters. (B) Quantified viability of HUVECs, hCMEC/D3, and MS1 encapsulated in microfibers with time (n=3). (C) Representative brightfield images of bioprinted multilayered lattice structures using μF_-cell_TP bioink. (D) Representative brightfield images of co-cultured bioprinted multilayered lattice structures using control, μF_-cell_TP and μF_+cell_TP bioinks with inclusion of laminin and VEGF. Scale bars: 200 μm (A and D) and 500 μm (C).

Then we formulated various μFTP bioinks for bioprinting practice using a photo-crosslinkable cell-laden precursor solution (5% GelMA and 2.5% gelatin solution carrying 5×10^6^/ml HUVECs) as the basic matrix phase. The relatively low concentration of GelMA (5%) was used to promote the growth of HUVECs and gelatin was added to enhance the printability, which could dissociate during 37 ℃ culturing. Sacrificial microfibers with or without HUVEC cells were mixed with the matrix phase solution at the mass ratio of 1:4 to formulate two μFTP bioinks, short for μF_-cell_TP and μF_+cell_TP, respectively. The microfiber-free group was taken as the bulk control. According to the previous optimization, 0.3% sodium citrate solution was chosen and supplemented to culture medium for the first 12 hours to dissolve the microfibers. According to the results of the Cell Counting Kit-8 assay (**Figure S8A**), μF_-cell_TP and μF_+cell_TP groups showed a higher metabolic activity than the bulk control, suggesting a better mass transfer condition with the introduction of channel pores, and the cells in the μF_+cell_TP group consistently showed the highest viability.

The brightfield and immunofluorescence staining of the bioprinted scaffolds (**Figure S8B** and **Figure S9**) showed that cells in the μF_+cell_TP group attached to the wall of channel pores and stretched from day 1 on, indicating the release and survival of in-situ delivered cells. In the μF_-cell_TP group, some cells still migrated to the channel pores and attached to the wall, though there was no cell loaded in the microfibers. Some cellular tube structures were seen inside the printed filaments both in the μF_-cell_TP and μF_+cell_TP groups from day 3 on, and the μF_+cell_TP group seemed to induce the formation of more confluent endothelium layers (**Figure S9**). In contrast, the number of ECs inside the filaments in the bulk control group decreased with time and most cells kept a round shape during the 7-day culture. The cells near the surface tended to grow and spread, forming a layer of cells on the outer surface on day 7. The migration and death of cells inside the hydrogel and confluent layer formation on the surface were also seen in literature when encapsulating HUVECs in bulk hydrogel filaments,[34] which, to some extent, promoted the formation of an endothelial layer on the outer surface. However, for our approach, this may also affect the nutrient acquisition of the inner cells. This is likely due to the tendency of ECs for physical interfaces and nutrients. We obtained similar results when we used hCMEC/D3 cells instead of HUVEC cells. On days 3 and 7, we observed tubular structures formed by cells adhering to the microchannel surfaces, which demonstrates the versatility of our μFTP strategy (**Figure S10**).

To further promote the growth of ECs and the formation of vasculatures, we continue the optimization of the μFTP formulations. Laminin is an important component of the basement membrane whose function is a mediator of cell adhesion to the matrix and binding to a variety of growth factors, such as vascular endothelial growth factor (VEGF).[35] Based on an optimized protocol in our lab, we incorporated 100 μg/ml laminin and 50 ng/ml VEGF into the matrix phase of the μFTP bioink. In addition, 2.5×10^6^/ml MRC-5 (human lung fibroblasts) were also supplemented to serve as a paracrine agent to support the growth of ECs. We printed multilayered lattice structures using updated bioink formulations and set up three experimental groups as before. Both the immunofluorescence and optical images (**Figure 6C-D**, **Figure 7A**) showed that HUVECs spread well inside the filaments and connected to each other. In the bulk control group, we observed some intercellular junctions within the first 3 days of culture. However, these connections nearly disappeared by day 7, and overall, the cells exhibited minimal spreading with no formation of tubular structures. In contrast, in both μFTP groups, the sacrificial microfibers provided a biological interface that offered sufficient space for cell adhesion and spreading. By day 3, abundant interwoven vessel-like tubular structures were observed within the bioprinted filaments, demonstrating the critical role of the μFTP strategy in promoting endothelialization. However, by day 7, the tubular structures in the μF_-cell_TP group had slightly faded, while the μF_+cell_TP group maintained a considerable endothelialization.

**Figure 7.**
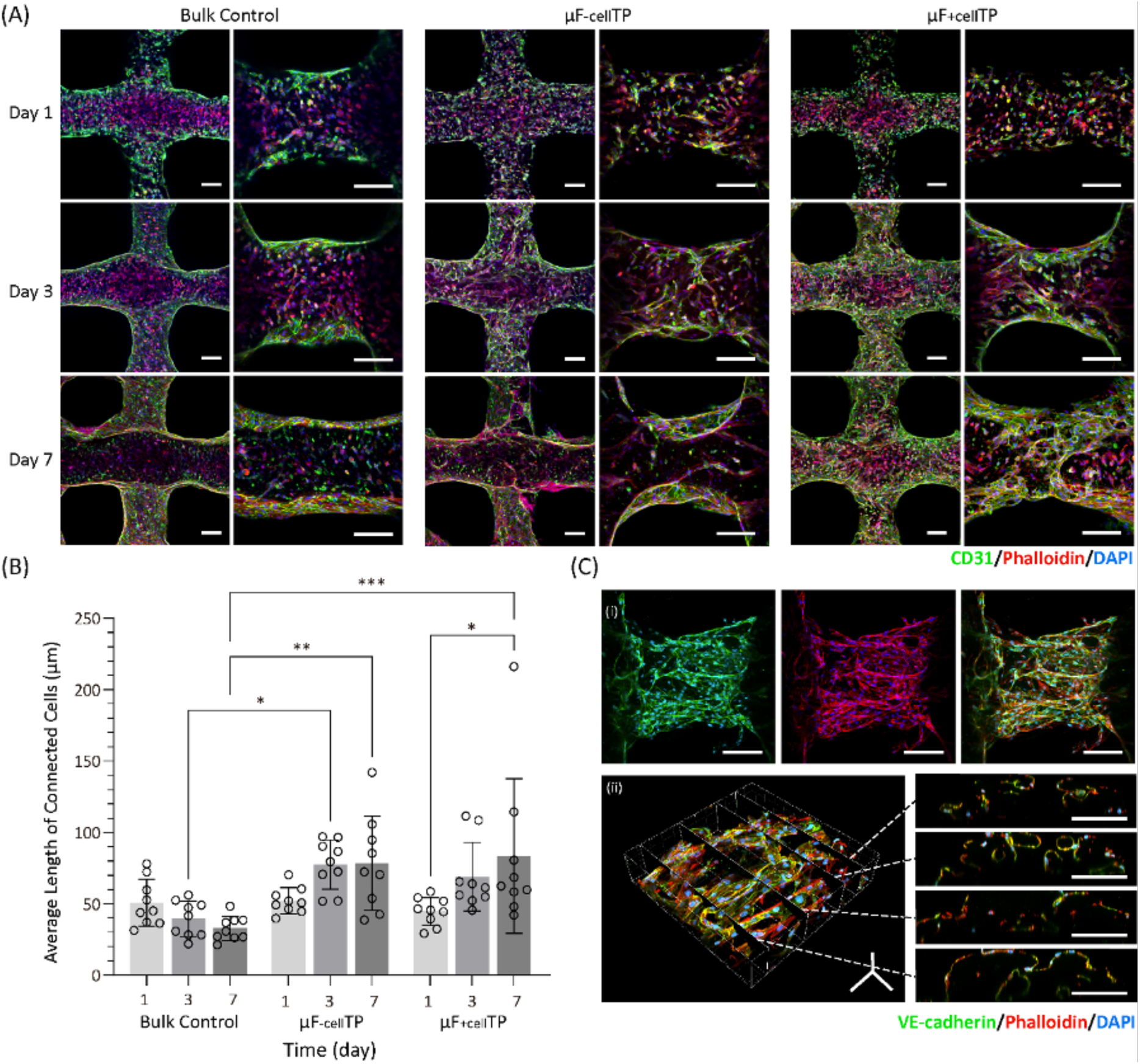
Microfiber-templated porogel bioink allows for spread of HUVECs within 3D printed structures. (A) Z-stack immunofluorescence images of co-culture 3D bioprinted structures with the inclusion of laminin and VEGF. CD31, F-actin, and DAPI are stained with green, red, and blue, respectively. (B) Quantification of average length of connected CD31 positive cells in 3D bioprinted structure with inclusion of laminin and VEGF (n=9). (C) (i) Z-stack immunofluorescence images of bioprinted filament and (ii) 3D reconstruction of immunofluorescence-stained scaffolds and cross-sectional view of the bioprinted filament of co-culture μF_+cell_TP group on day 7. VE-cadherin, F-actin, and DAPI are stained with green, red, and blue, respectively. Scale bars: 200 μm (A and C-i), and 100 μm (E-ii). One-way ANOVA with a Tukey’s multiple comparisons test, *P < 0.05; **P < 0.01; ***P < 0.001; ns, not significant.

We quantified the growth of endothelial cells within bioprinted filaments by calculating the length of CD31 positive region with ImageJ, which indicates the connection level of endothelial cells (**Figure S11, Figure 7B**). Slices at different z-axis depths from the confocal images were selected for statistical analysis. We applied a series of image processing steps to the original images, ultimately obtaining lines that represent connected endothelial cells, which enabled the calculation of their lengths. The length values in the control group gradually declined over time, reaching the lowest among all groups by day 7. This decrease is probably attributed to the lack of space for cell spreading and the limited nutrient supply within the printed filaments, which lead to a large amount migration of cells from the interior to the surface or cell death. Similar to the results shown in the fluorescence images, the μFTP groups exhibited a great increasing trend in endothelialization during seven days. The length of connected cells in μF_+cell_TP group showed a higher increase trend by day 7 compared with μF_-cell_TP group, reaching the highest value among all groups. This outcome may be because, compared to the μF_-cell_TP group, endothelial cells in the microchannels of the μF_+cell_TP group are more concentrated, which enhances tight junctions between endothelial cells and leads to closer cell-cell interactions, helping to maintain endothelialization. Additionally, compared to the μF_-cell_TP group, the μF_+cell_TP group exhibited a higher initial cell density within the microchannels. Given the substantial nutrient consumption inside the printed filaments, there were more cells migrating outward from the internal pores, which enhances the connectivity between the inner and outer interfaces and promotes nutrient transport.

We further stained VE-cadherin, an endothelial-specific adhesion molecule, to show tight junctions between cells and the formation of the endothelial barrier (**Figure 7C-i**). Even though the cells were relatively dispersed on the first day, they interconnected within the channels to form well-defined tubular structures, and stronger VE-cadherin expression at the cell boundaries was observed on days 3 and 7, indicating the formation of tight intercellular junctions. ICAM-1 (Intercellular Adhesion Molecule-1), which is typically upregulated during inflammatory responses and facilitates the adhesion of leukocytes to endothelial cells, was observed to be expressed in bioprinted μF_+cell_TP structures (**Figure S12**). The secretion of von Willebrand factor (vWF) was found in the bioprinted structure of the μF_+cell_TP group, which plays a crucial role in thrombogenicity and could further support the maintenance of endothelial cell function (**Figure S12**). The higher magnification immunofluorescent images and 3D reconstruction image of μF_+cell_TP group clearly presented the 3D tubular structures within the filament (**Figure 7C-ii, Figure S13**). The cross-sectional view of the filament clearly shows that HUVECs attached to the microchannel densely in all directions and showed a circular cross-section. Such densely formed vessel-like structures inside the printed filaments were not random but universally occurred throughout the whole structures, evidenced by the panorama imaging of the whole scaffold (**Figure S14**). Compared to previous work based on the microfiber-template method, we have successfully transferred the methodology to 3D bioprinting and achieved a higher density of endothelialized tubular structures within the hydrogel due to the optimization of the ink components. Moreover, this density is adjustable. Additionally, we performed paraffin slice and H&E staining on samples from the μF_+cell_TP groups at different time points to achieve clearer characterization of the internal structure of the printed constructs (**Figure S15**). The staining results revealed many pores within the printed structures, with cells growing inside. In certain areas, cells were observed adhering to the pore surfaces in cross-sections, forming ring-like patterns.

In conclusion, we engineer tubular interfaces down to the building blocks of 3D bioprinting for enhanced microvascularization, which has never been reported before. Although the general porogen-dispersing approach is limited in controlling the position and direction of each porogen, our μFTP strategy represents a way to address the enduring trade-off between fabrication resolution and throughput for vascular engineering. Specifically, we develop the μFTP bioink system and incorporate an in-situ endothelialization approach, where the microfibers act as both porogen and carriers for ECs in the 3D bioprinting scenario. Our optimization (e.g., microfiber softening and in-situ cell seeding) allows us to print customized constructs with a wide range of proportions of tubular porosity and endothelialization free of post-cell seeding.

### 2.5. *In vivo* vascularization

To further investigate whether the μFTP system could promote the *in vivo* ingrowth of tissues and blood vessels, we bioprinted scaffolds with or without microchannels and implanted them subcutaneously into 4-week-old male SD rats for 4 weeks. To favor the biocompatibility of scaffolds and vascular invasion capability, we selected 7.5% GelMA supplemented with 100 μg/mL rat laminin and 50 ng/mL rat VEGF as the basic matrix material. Three groups of cubic scaffolds (10mm×10mm×1.2mm) were 3D printed, including the bulk control group (no microfiber added in the matrix), low porosity μFTP group (1:4 microfiber to matrix ratio), and high porosity μFTP group (1:2 microfiber to matrix ratio). The microfibers were properly removed before implantation and the scaffolds were taken out and characterized at week 2 and week 4. The H&E staining results indicated a clear gel area in the control group, suggesting a minimal growth of the native tissues (**Figure 8A**). In contrast, a large number of cells and tissues were found in the scaffold microchannels in the μFTP groups at both week 2 and week 4, showing a significant tissue in-growth (**Figure 8B**). At week 4, the cells and tissues occupied almost all the pores and fused to the scaffolds firmly. The Masson staining results further confirmed the conclusions above and showed a thick collagen layer in the tissue-scaffold interfaces in the bulk control group, while the μFTP groups exhibit uniform collagen both in the interfaces and inside the bioprinted constructs. The immunohistochemistry staining of CD31 demonstrated obvious CD31 expression in the tissues that grew into the scaffolds in the μFTP groups and CD31-expressing cells formed lumens that indicate the location of blood vessels. This demonstrates the ability of the scaffold to induce the vessel in-growth, showing a promising in vivo vascularization (**Figure 8C**). To further verify the maturation of blood vessels growing into the interior of the hydrogel, we conducted the immunohistochemical staining for VE-cadherin, VEGF, and Nestin (**Figure S16**). In the μFTP groups in week 2 and week 4, clear VE-cadherin-expressing was observed in tissues that grew into the voids of the hydrogel, which proved the presence of blood vessels and integrity of barrier functions. For VEGF, dark regions were found in abundance at the hydrogel-host tissue interface in bulk control, while VEGF expression was massively presented in the implants together with the infiltrated tissues in the μFTP groups. Additionally, noticeable Nestin expression was observed within the hydrogel in the μFTP groups in week 2, indicating the presence of neural progenitor cells and suggesting the possibility of ongoing angiogenesis. However, by week 4, Nestin expression was not as prominent as in week 2, which might indicate the in-growth tissues and vessels tend to be mature.

**Figure 8.**
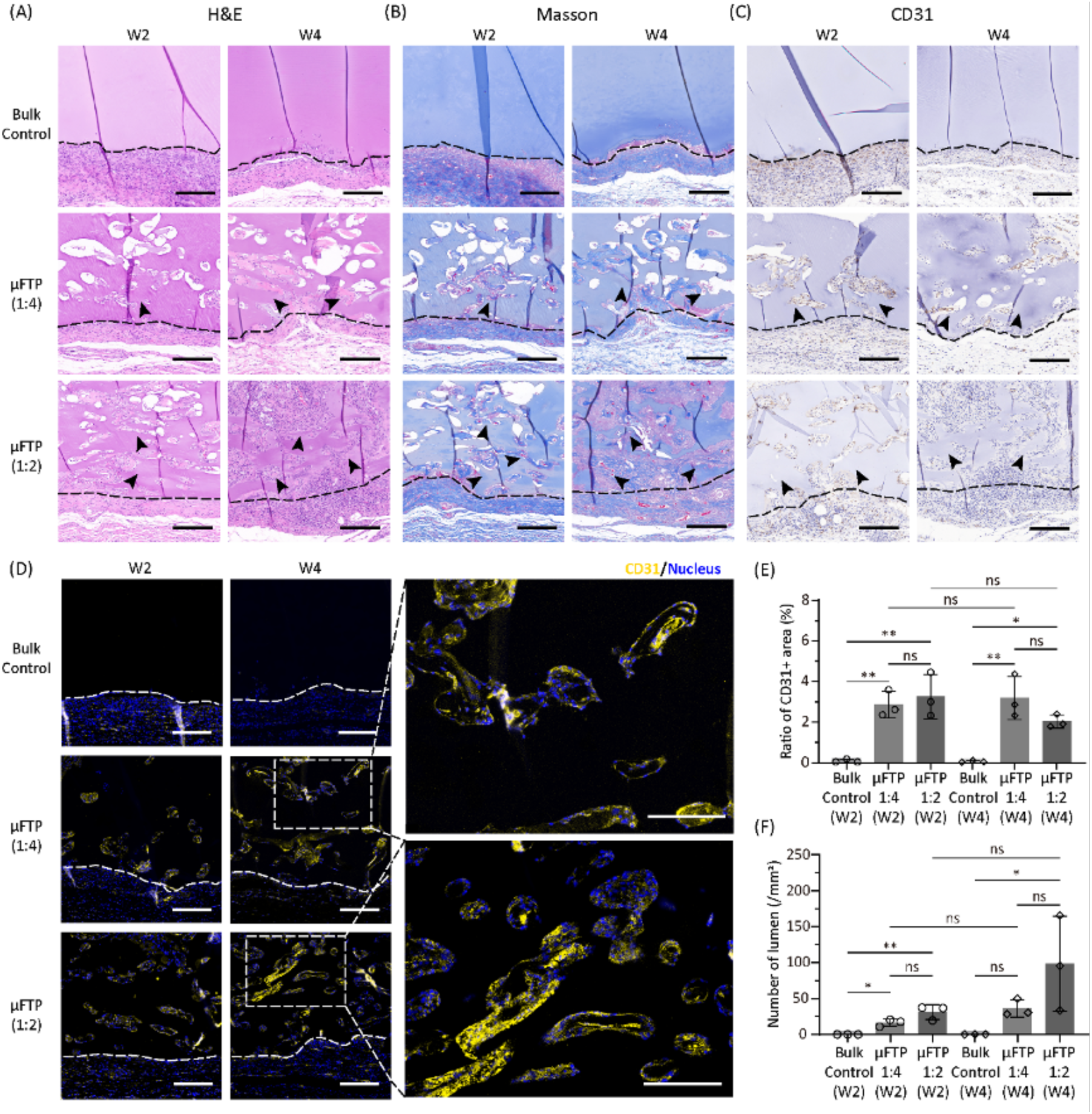
Microfiber-templated porogel enhances in-growth of native tissues and vessels in rat subcutaneous models. Representative microscopic images of (A) H&E staining, (B) Masson staining, (C) immunohistochemistry staining of CD31 (the dot line indicates hydrogel interface with animal tissue and the arrow shows tissue in-growth) and (D) immunofluorescence staining of CD31 of 3D printed structures at week 2 and week 4 post-implantation. (E) Quantification of CD31+ area inside the implants. (F) Quantification of the number of CD31+ lumen inside the implants. Scale bars: 200 μm (A-D). One-way ANOVA with a Tukey’s multiple comparisons test, *P < 0.05; **P < 0.01; ***P < 0.001; ns, not significant (n = 3).

Immunofluorescence staining of CD31 was conducted to further visualize the vessels (**Figure 8D**). Aligning with the above results, CD31 positive area and mature luminal structures were seen only inside the μFTP scaffolds but not the bulk control. Compared to the result on week 2, the results on week 4 showed a larger number of CD31-positive regions with stronger fluorescence expression. The hollow structures outlined by the fluorescence were also more clearly defined. The quantification results further confirm the significant differences in both CD31 positive areas and lumen numbers (**Figure 8E-F**). Regarding the comparison of porosity, the CD31 positive area was higher in 1:2 group than that in 1:4 group at week 2, while more CD31 positive area was seen in the 1:4 group at week 4. Meanwhile, 1:2 group showed more lumen numbers throughout the four weeks. These results suggest that the scaffolds with high porosity might induce the growth of more capillaries rather than larger vessels after four weeks, which demonstrated a better integration between implanted scaffold and native tissues.

To further compare the vascularization potential of μFTP hydrogels and conventional microgel-templated hydrogels *in vivo*, we conducted a subcutaneous implantation study comparing bioprinted microgel-templated group and μFTP group with a fixed porogen-to-matrix ratio of 1:4, together with a bulk control group (**Figure S17**). Macroscopic images showed that the bulk group exhibited no significant vascular ingrowth or attachment. Compared to the microgel-templated group, the μFTP group had significantly more surrounding vasculatures inside. Subsequently, the hydrogel samples were extracted, and the adhered host tissue was removed as much as possible. Under the microscope, vascular ingrowth was observed in both porous hydrogel groups. In the μFTP group, the vessels at the sample edges exhibited a clear directional orientation with smaller diameters. This is likely due to the microfiber orientation formed during the bioprinting process, which guided host vessels into the microchannels. In contrast, no significant directional vascularization was observed in the microgel-templated group. Previous studies have confirmed the phenomenon of host blood vessels entering the microchannels within implants and integrating with the host circulatory system.[25] This suggests the possibility of directionally customized growth and regeneration of host blood vessels induced by the microchannels aligned with designed printing path, which needs further investigation in the future. To confirm the connectivity and perfusability between the ingrown vessels within the hydrogel and the host vasculature, FITC-Dextran was administered through the tail vein (**Figure S18**). After allowing time for circulation, hydrogel implants were taken out with the adhered host tissue removed as thoroughly as possible. Under fluorescence microscopy, green fluorescence was observed in the vessels within the μFTP hydrogel, demonstrating that FITC-dextran had reached the implantation position through the circulatory system. In conclusion, our results have demonstrated that the μFTP inks could significantly promote tissue and blood vessel in-growth *in vivo*.

## 3. Conclusion

In summary, based on the sacrificial material-aided *in situ* cell delivery strategy, we developed a microfiber-templated porogel (μFTP) bioink system to fabricate customized 3D structure incorporated with tunable microchannels and thus vasculatures. The μFTP bioink exhibits good printability and enables the construction of complex, large-scale structures with varied porosities (up to 55%) and void channel sizes (from ∼50 to ∼150 μm). *In vitro* experiments demonstrate that a faster and higher degree of endothelialization could be achieved by *in situ* cell delivery within the microchannels. The incorporation of biochemical cues further enhances the cellular microenvironment of μFTP formulation and induces tightly connected endothelium layer. *In vivo* experiments also evidence that the introduction of microchannels in bioprinted implants leads to the promotion of the in-growth of blood vessels and native tissues compared to the bulk counterpart under the same matrix material condition. Overall, the μFTP bioink provides a general strategy of microchannel formation down to the building block level and allows for *in-situ* endothelialization of microvasculatures, facilitating vascularization of engineered *in vitro* tissues.

## Author Contributions

Y. Guo and L.Ouyang designed the study. Y. Guo performed the experiments. Y. Guo and L.Ouyang wrote the manuscript. L.Ouyang supervised the study. Y. Guo and Z. Wang contributed to animal experiments. X. Zhang contributed to the co-axial needles and microfibers preparation, cell culture and immunofluorescence staining. J. Li contributed to the microfiber preparation and fiber dissolution characterization. Y. Guo and X. Zhang contributed to the figure design and drawing. L. Ouyang gained the funding.

## Acknowledgements

The authors acknowledge R. Xu and B. Dou from the Ouyang Group for kindly help with cell culture, D. Zhou and C. Hua from the Ouyang Group for kindly help with material preparation and experiment design, Y. Lv and S. Gao from Peking University Third Hospital for initial discussion on animal experiments. L.O. acknowledges the financial support from National Natural Science Foundation of China (No. 52105306, No. 52475305, No. 32211530075) and Faculty Startup Funding from Tsinghua University (012-533302004).

## Data Availability Statement

All data needed to evaluate the conclusions in the paper are present in the paper and/or the Supplementary Materials.

## Ethical Statement

The authors declare no ethical issues.

